# Susceptibility of Spotted Doves (*Streptopelia chinensis*) to Experimental Infection with the SFTS Phlebovirus

**DOI:** 10.1101/466094

**Authors:** Zhifeng Li, Changjun Bao, Jianli Hu, Chengfeng Gao, Nan Zhang, Huo Xiang, Carol J. Cardona, Zheng Xing

## Abstract

**Background:** Severe fever with thrombocytopenia syndrome virus (SFTSV), an emerging human pathogen naturally transmitted by ticks, has spread widely during the last few years. Although SFTSV has been detected in wild birds, the natural reservoir and amplifying hosts for the virus have not been well-studied.

**Methodology/Principle Findings:** Here we report an experimental infection of spotted doves (*Streptopelia chinensis*) with two strains of SFTSV, JS2010-14 (hereafter JS2010), a Chinese lineage strain and JS2014-16 (JS2014) from a Japanese lineage, which represent the main viral genotypes currently circulating in East Asia. We determined that spotted doves were susceptible to SFTSV and the severity of the viremia was dose-dependent. When challenged with 10^7^ and 10^5^ PFU, all doves developed viremia which peaked 3-5 days post-infection (dpi). A subset (25-62.5%) of the birds challenged at 10^3^ PFU, developed viremia. Virulence of SFTSV in spotted doves appeared to be strain-dependent. Infection with the strain of JS2014 led to a death rate of 12.5% and higher viremia titers in experimentally inoculated birds. The doves inoculated with the JS2010 strain survived infection with relatively lower virus titers in the blood.

**Conclusions/Significance:** Our results suggest that spotted doves, one of the most abundant bird species in China, could be a competent amplifying host of SFTSV, the strain of the Japanese lineage in particular, with higher viremia titers and play an important role in the transmission of SFTSV. Our observations shed light on the ecology of SFTSV which could benefit the implementation of future surveillance and control programs.

**Author Summary:** Severe fever with thrombocytopenia syndrome virus (SFTSV), an emerging human pathogen naturally transmitted by ticks. Our recent study have showed that some species of migratory birds, such as swan geese and spotted doves, could be parasitized by *H. longicornis*, and antibodies against the virus could also be determined in these birds, which showed that migratory birds could be infected by SFTSV naturally. Other studies have reported that migratory bird routes and the distribution of *H. longicornis* in East Asia overlap with the geographic distribution of SFTSV. Migratory birds are known to be carriers and transmitters of infectious agents, like the causative agents of influenza, West Nile encephalitis, and Lyme disease. Wild birds often travel long distances carrying various parasites, including ticks, which may be infected with viruses and bacteria. It is therefore reasonable to hypothesize that migratory birds may have played an important role in dispersing *H. longicornis*-borne SFTSV in both scenarios, either the birds are infected directly with the virus or the birds are carriers of parasitic ticks that are infected with the virus. Here we report an experimental infection of spotted doves (*Streptopelia chinensis*) with two strains of SFTSV, JS2010-14 (hereafter JS2010), a Chinese lineage strain and JS2014-16 (JS2014) from a Japanese lineage, which represent the main viral genotypes currently circulating in East Asia. We determined that spotted doves were susceptible to SFTSV and the severity of the viremia was dose-dependent.

Interestingly, virulence of SFTSV in spotted doves appeared to be strain-dependent. Infection with the strain of JS2014 led to a death rate of 12.5% and higher viremia titers in experimentally inoculated birds. The doves inoculated with the JS2010 strain survived infection with relatively lower virus titers in the blood. These findings provide novel insights for understanding the rapid spread of the virus in a short time span, especially the SFTSV strains from the Japanese lineage (genotype E), which presented cross ocean transmission.

## Introduction

Severe fever with thrombocytopenia syndrome virus (SFTSV) is a *phlebovirus* in the family *Phenuiviridae* and causes severe fever with thrombocytopenia syndrome (SFTS), a severe hemorrhagic fever disease in East Asia (1, 2). The disease is characterized by high fever and a drastic reduction of platelets and leukocytes resulting in multi-organ failure with mortality up to 10% of infected humans. SFTSV was firstly isolated from a patient in Eastern China in 2010. By the end of 2017, more than 12,000 cases have been reported in 23 Chinese provinces and are why SFTS has become an increasingly important public health concern (3-6).

Severe Fever with Thrombocytopenia virus is a tick-borne zoonotic virus that has been detected in or isolated from several species of ticks, especially *H. longicornis*, a widely distributed tick species in East Asia [7-9]. SFTSV has a broad spectrum of animal hosts. Previous studies conducted in East Aisa including China, South Korea, and Japan showed that many domestic and wild animals were susceptible to SFTSV infection resulting no or inconspicuous clinical signs [10-13]. Additionally, our study showed that some species of migratory birds, such as swan goose (*Anser cygnoides*) and spotted doves, could be parasitized by *H. longicornis* and infected by SFTSV, which might contribute to a long-distance spread of SFTSV via migratory flyways [7]. This hypothesis could explain why SFTSV has spread rapidly in China and genetically-close viral strains were identified both in China and Japan or Korea within a relatively short time span in the past years. Experimental infection with SFTSV could result in mild signs with a moderate viremia levels in vertebrate animals, which might serve as amplifying hosts in the natural transmission cycle of SFTSV [14-16]. However, how susceptible avian species could be to SFTSV has not been well studied.

Spotted doves are a common migratory bird in China. This species is found in most parts of China in summer months, but in winter, most migrate to warmer areas of southern China [28]. In this study, we challenged naive spotted doves with two genotypes of SFTSV to establish an avian model of infection. Our objective was to determine the susceptibility of spotted doves to SFTSV infection, examine its virulence and duration of viremia, to assess the potential role of doves as a competent host capable of transmitting the virus.

## Materials and Methods

### Ethics Statement

All shipment of birds, daily husbandry, and study protocols were handled in strict accordance with the Animal Ethics Procedures and Guidelines of the People’s Republic of China (Regulations for Administration of Affairs Concerning Experimental Animals, China, 1988), and were pre-reviewed and approved by the Ethics Committee of the Jiangsu Provincial Center for Disease Control and Prevention (Certificate No. JSCDCLL [2016]032). All birds used in this study were euthanized under isoflurane anesthesia.

### Sources of viruses and birds

Two SFTSV strains, JS2010-14 of Chinese (hereafter JS2010) and JS2014-16 (hereafter JS2014) of Japanese lineages, were used in the study. These two viral strains were isolated from local confirmed SFTS cases in 2010 and 2014, respectively. The spotted doves were purchased from a commercial breeder of the species in China and held for 2 weeks in observation prior to SFTSV challenge. The birds used for the study were determined to be clinically normal by a qualified veterinarian. Upon arrival each bird was given a numbered leg band, and caged in Biosafety Level 3 Animal facilities. The spotted doves were provided 12 hr light/12 hr darkness housed in groups of six of the same gender in wire cages measuring approximately 80 cm (long) x 60 cm (wide) x 60 cm (height), and were provided a commercial seed mix and water. All enrolled birds were males, 2 months of age.

### Challenge of spotted doves

Two independent challenge studies were conducted.

In the first, the birds were randomly assigned to four treatment groups for each SFTSV strain: procedural controls (n = 4), and three SFTSV challenge groups, each given a different SFTSV dose: 10^3^ (n = 8), 10^5^ (n = 8), and 10^7^ (n = 8) PFU (Table 1). The birds in the control group were housed together separately from the virus challenged groups. On day 0 of the study, control birds were injected subcutaneously (s.c.) in the medial left thigh with 100 μL of serum-free Dulbecco’s Modified Eagle Medium (DMEM) as previously described [17]. The individual birds in the three SFTSV challenge groups were each inoculated s.c. with 100 μL of DMEM containing 10^7^, 10^5^ or 10^3^ PFU of a low passage (<3) human origin isolate of SFTSV according to their group assignment (Table 1). A bird was considered infected with SFTSV if live virus was isolated from a serum sample at any sampling time point, or if the bird developed anti-SFTSV antibodies. Each of the birds in the virus-inoculated groups was sampled on day 1 through 14 post-inoculation (pi). On each sampling day, a 100 μL of blood was collected, Whole blood was allowed to clot for 30 min at room temperature in blood collection tubes and held at 4°C until centrifugation at 2000 x g for 10 min. Sera were collected and diluted in DMEM for a final serum:media dilution of 1:5. The resulting diluted serum samples were stored at −80°C until testing.

**Table 1.**
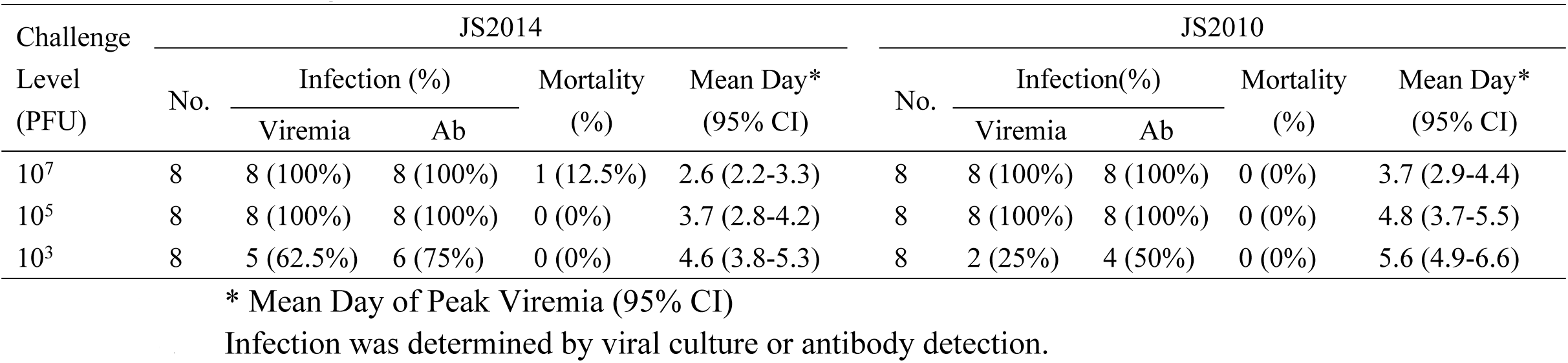
Spotted Doves susceptibility to SFTSV infection is dependent on challenge level.

| Challenge Level (PFU) | JS2014 |  |  |  |  | JS2010 |  |  |  |  |
| --- | --- | --- | --- | --- | --- | --- | --- | --- | --- | --- |
|  | No. | Infection (%) |  | Mortality (%) | Mean Day* (95% CI) | No. | Infection(%) |  | Mortality (%) | Mean Day* (95% CI) |
|  |  | Viremia | Ab |  |  |  | Viremia | Ab |  |  |
| 10<sup>7</sup> | 8 | 8 (100%) | 8 (100%) | 1 (12.5%) | 2.6 (2.2-3.3) | 8 | 8 (100%) | 8 (100%) | 0 (0%) | 3.7 (2.9-4.4) |
| 10<sup>5</sup> | 8 | 8 (100%) | 8 (100%) | 0 (0%) | 3.7 (2.8-4.2) | 8 | 8 (100%) | 8 (100%) | 0 (0%) | 4.8 (3.7-5.5) |
| 10<sup>3</sup> | 8 | 5 (62.5%) | 6 (75%) | 0 (0%) | 4.6 (3.8-5.3) | 8 | 2 (25%) | 4 (50%) | 0 (0%) | 5.6 (4.9-6.6) |

\* Mean Day of Peak Viremia (95% CI) Infection was determined by viral culture or antibody detection.

Following SFTSV challenge, birds were observed daily for clinical signs for 14 dpi. Birds with difficulty perching or other neurological signs, or that were moribund were humanely euthanized. At 14 dpi all birds were euthanized under isoflurane anesthesia.

In the second study, for each of the SFTSV strains, 16 spotted doves were inoculated with a dose of 10^5^ PFU with an additional two birds inoculated with serum-free DMEM to serve as negative controls, respectively,. Birds were tested for infection with SFTSV by viral isolation from organs and tissues in addition to serum and/or antibody detection as in the previous experiment. Three randomly selected birds were sacrificed at 2, 4, 7, and 14 dpi for necropsy and heart, liver, lung, spleen, kidney, and brain collected. A small portion of each organ was sub-sampled, weighed, and homogenized in 1 mL of DMEM containing 100 μg/mL of penicillin and streptomycin using a mini-bead beater instrument (TissueLyser LT, Qiagen, Germany) for virus isolation performed in Vero cell culture and confirmed with real-time RT-PCR. The negative control birds were sacrificed and necropsied at 14 dpi, and their organs processed as described. The serum samples were tested for the presence of anti-SFTSV antibodies at 2, 4, 7, and 14 dpi. Surviving birds were euthanized and necropsied at 14 dpi.

### Virus isolation and titration

Virus titration was performed as described [17] using a 24-well plate for a mini-plaque technique to accommodate small sample volumes. 0.5 ml of Vero cells (2 × 10^5^ cells/ml in stock) in DMEM media was added to each well. The plates were incubated for 4 days at 37°C in an incubator with 5% CO_2_. Sera and homogenized organ samples were individually centrifuged at low speed for clarification. The supernatants were diluted at 1:10 in DMEM medium containing 10% fetal bovine serum. An individual diluted viral inoculum was added to three wells containing confluent Vero cell monolayers in the 24-well plate and in incubated for 45 min after which it was removed. 1 ml of complete agarose overlay was added to each well. Plaques formed in the positive wells were further titrated on six-well plates in a ten-fold dilution series until a countable endpoint was reached. Cell cultures were checked for plaque formation at 96, 120, and 144 hrs pi and the number of plaques was recorded. Infectious virus titers were calculated as PFU/ml or PFU/cm^3^ tissue.

### Virus detection by RT-PCR

RNA was extracted from 140 μL of diluted serum samples in serum-free DMEM using the QIAamp Viral RNA Mini kit (Qiagen). RNA was extracted from the brain using the RNeasy Lipid Tissue Mini extraction kits (Qiagen), and from the remaining tissues using the RNeasy Mini extraction kit (Qiagen). Real-time RT-PCR was performed using the QuantiTech RT-PCR kit (Qiagen). The primers were designed as previously described and used in a one-step real-time RT-PCR [18]. The forward (S-for)/reverse (S-rev) primers and MGB probe (S-pro) used in the real-time RT-PCR were targeted to the S segment of the viral genome. Conditions for the reaction were as follows: 50°C for 30 min, 95°C for 15 min, 40 cycles at 95°C for 15 sec, and 60°C for 1 min. Amplification and detection were performed with an Applied Biosystems 7500 Real-time PCR system (Applied Biosystems, Foster City, CA). Data were analyzed using the software supplied by the manufacturer.

### Antibody detection

All serum samples were heat-inactivated at 56°C for 30 min and tested for anti-SFTSV antibodies by plaque reduction neutralization assay (PRNT) on 12-well plates as described [15]. Samples exhibiting a neutralization of ≧90% were considered positive for antibodies to SFTSV (PRNT_90_). Additional sera were tested for both IgG and IgM SFTSV antibodies with a commercial double antigen sandwich ELISA kit from Xinlianxin Biotech (Wuxi, China). Positive sera were 2-fold diluted starting at 1:10 for the assay to obtain endpoint titers determined by the cutoff values set by the positive and negative ELISA controls.

### Statistical analysis

All statistical analyses were performed with SPSS 19.0 (SPSS, Chicago, IL) and statistical significance level was set at 0.05. For categorical data, the proportion and 95% confidence interval (CI) were calculated and differences in proportions were compared with the Fisher’s exact test. Unless indicated, all tests of proportions or means were two-sided.

## Results

### Susceptibility of Spotted Doves to SFTSV

To investigate the susceptibility of spotted doves to SFTSV infection, the birds were grouped into four groups and each group challenged with the two SFTSV strains at doses of 10^7^, 10^5^ and 10^3^ PFU, respectively. The last group was inoculated with the serum-free DMEM vehicle control. Our data demonstrates that spotted doves were infected and developed viremia with both viral strains (Table 1, Figure 1). Viremia appeared in each of the birds challenged with the doses of 10^7^ and 10^5^ PFU of both viral strains. When challenged with the dose of 10^3^ PFU, viremia was detected in fewer birds than in the groups challenged with the higher doses. On the other hand, when challenged at 10^3^ PFU, more birds were viremic after challenge with strain JS2014 of the Japanese lineage (5/8) than with strain JS2010 of the Chinese lineage (2/8) (Table 1).

**Figure 1.**
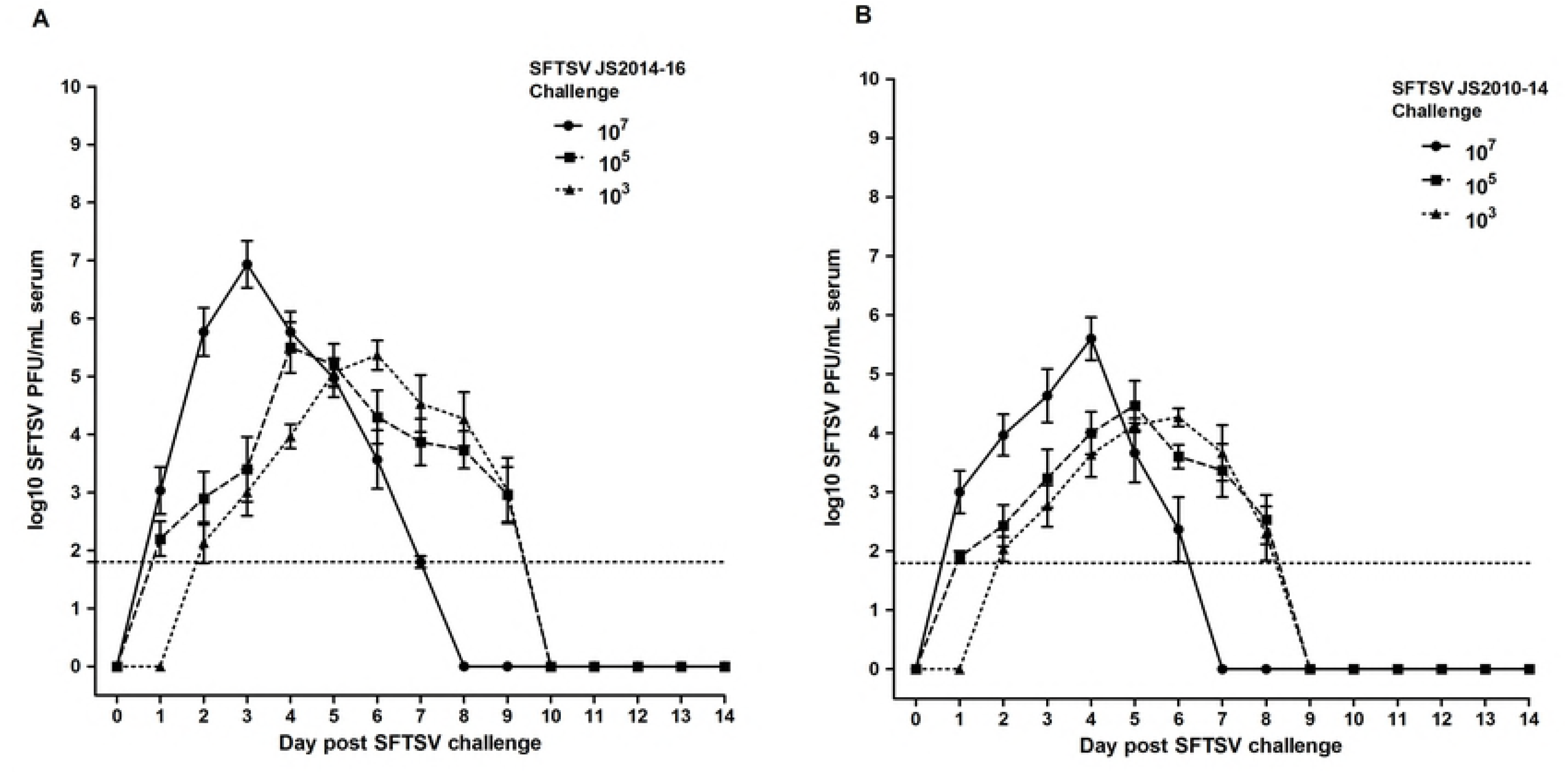
Experimental infection of Spotted Doves with two SFTSV strains. Error bars represent standard error of the mean log_10_ PFU/mL serum. The horizontal dashed line indicates a limit of detection of 10^1.8^ PFU/mL. A: Spotted Doves inoculated with 10^7^ plaque forming units (PFU) of SFTSV strain JS2014 (solid line) had higher mean viremia and earlier peak viremia than birds inoculated with 10^5^ (dashed line) or 10^3^ PFU (fine dashed line) of virus. B: Spotted Doves inoculated with 10^7^ plaque forming units (PFU) of SFTSV strain JS2010 (solid line) had higher mean viremia and earlier peak viremia than birds inoculated with 10^5^ (dashed line) or 10^3^ PFU (fine dashed line) of virus.

We were able to detect SFTSV-specific antibodies in doves challenged with each of the SFTSV strains. In the 10^3^ PFU group with JS2010 and JS2014 SFTSV strains, two and one additional birds developed anti-SFTSV antibodies without having been viremic, respectively. All control birds (n=4) were negative by viral isolation, viral specific antibodies, and viral RNA by real-time RT-PCR and none died.

Mortality was observed only in the group of the birds challenged at 10^7^ PFU with the strain JS2014 (1/8, 12.5%) on day 7 pi. No birds died in the groups inoculated with 10^5^ and 10^3^ PFU of either virus strain and with no birds given the JS2010 strain at 10^7^ PFU. The data suggest that the infection of SFTSV in spotted doves was primarily self-restricted and infected birds recovered after a defined period of viremia.

### Dynamics of viremia in spotted doves challenged with SFTSV

Mean SFTSV viremia levels peaked on 3 dpi in the 10^7^ PFU challenge group of SFTSV strain JS2014 (mean=10^6.9^ PFU/mL, SD 10^0.3^), followed by the 10^5^ PFU group on 4 dpi (mean = 10^5.5^ PFU/mL, SD 10^0.2^), and by the 10^3^ PFU group on day 5 (mean = 10^5.3^ PFU/mL, SD 10^0.2^). The Mean SFTSV viremia of 10^7^, 10^5^, and 10^3^ PFU challenge groups of SFTSV strain JS2010 peaked on 4 dpi (mean=10^5.6^ PFU/mL, SD 10^0.3^), 5 dpi (mean = 10^4.5^ PFU/mL, SD 10^0.4^), and 6 dpi (mean=10^4.3^ PFU/mL, SD 10^0.2^), respectively. The mean peak day of viremia for individual birds of the 10^7^ PFU challenge group of SFTSV strain JS2014 (mean 2.6 d) occurred significantly earlier as compared to the other challenge groups (Table 1) (overall F=9.2, p<0.01, Tukey’s multiple comparisons of 10^7^ mean to 10^5^ and 10^3^ PFU, q=5.2 and q=5.1, respectively). With SFTSV strain JS2010, the mean peak day of viremia for individual birds of the 10^7^ PFU challenge group was 3.7 d and was also significantly earlier compared to the other challenge groups (Table 1) (overall F=10.2, p<0.01, Tukey’s multiple comparisons of 10^7^ mean to 10^5^ and 10^3^ PFU, q=6.2 and q=5.5, respectively). Viremia detected in birds of the 10^7^ PFU challenge group of two SFTSV strains also fell to the threshold of detection (5-6 days) more rapidly than the 10^5^ or 10^3^ PFU challenge groups (7-8 days).

### Multi-organ tropism of SFTSV in spotted doves

In the second study, the virus was successfully cultured from sera or organs in 15 of the 16 spotted doves challenged with 10^5^ PFU of SFTSV strain JS2014. Additionally, anti-SFTSV antibodies were detected in the final bird demonstrating that all of the birds were infected. Mortality was not observed. While of the10^5^ PFU JS2010 SFTSV challenge group, 12 of the 16 spotted doves were positive by viral isolation and two additional birds developed anti-SFTSV antibodies.

In the second study in the birds challenged with 10^5^ PFU of JS2014, SFTSV was detected through viral isolation or RT-PCR in multiple organs, including kidney, liver, heart, lung, and spleen taken from three sacrificed birds at 2 dpi, 4 dpi and 7 dpi (Table 2). At 14 dpi, however, SFTSV was no longer detectable by either viral isolation or RT-PCR in any tissue obtained from three sacrificed birds. For the birds inoculated with 10^5^ PFU of strain JS2010, SFTSV was detected by RT-PCR only in the spleen of the three sacrificed birds at 2 dpi. At 4 and 7 dpi, kidney, liver, heart, lung, and spleen were all positive with either viral isolation or RT-PCR in all three sacrificed birds. At 14 dpi, SFTSV was not detected by either method in any tissues from the sacrificed birds. (Figure 2).

**Figure 2.**
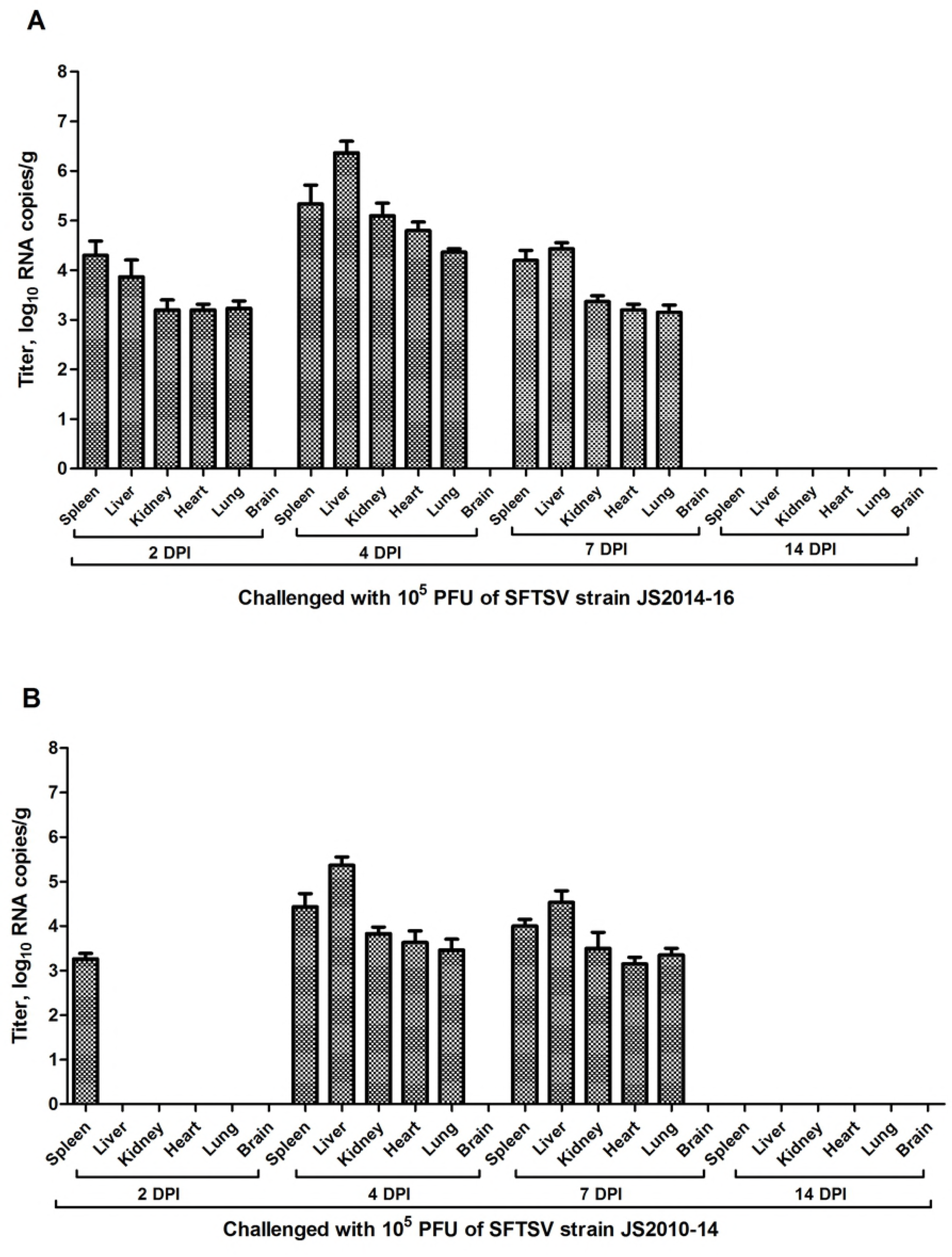
Tissue viral load, as determined by RNA copy numbers, in organs harvested from birds experimentally infected with 10^5^ PFU of two SFTSV strains on day different dpi. Viral titers are represented as geometric mean±SD. A detection limit of 0.95 log_10_RNA copies g^−1^ was determined. Spotted Doves inoculated with 10^5^ PFU of SFTSV strain JS2014 (A) had higher and earlier mean viremia than birds inoculated with 10^5^ PFU of SFTSV strain JS2010 (B).

### Development of SFTSV antibodies in spotted doves

Prior to the study we sampled the blood of all birds that were seronegative for specific antibodies to SFTSV by the PRNT assay. Final serum samples were collected at 14 dpi or at the time of death in the first study, and the antibodies to SFTSV were detected by the PRNT_90_ in 8/8 (100%), 8/8 (100%), and 6/8 (75%) of the SFTSV infected birds in the groups challenged with 10^7^, 10^5^, and 10^3^ PFU of the strain JS2014, respectively. In the groups challenged with 10^7^, 10^5^, and 10^3^ PFU of the strain JS2010, specific antibodies to SFTSV were detected in 8/8 (100%), 8/8 (100%), and 4/8 (50%) of the infected birds, respectively (Table 1). Neutralized antibodies for SFTSV were detected earlier in the spotted doves challenged with 10^7^ and 10^5^ PFU than in birds given 10^3^ PFU (Figure 3).

**Figure 3.**
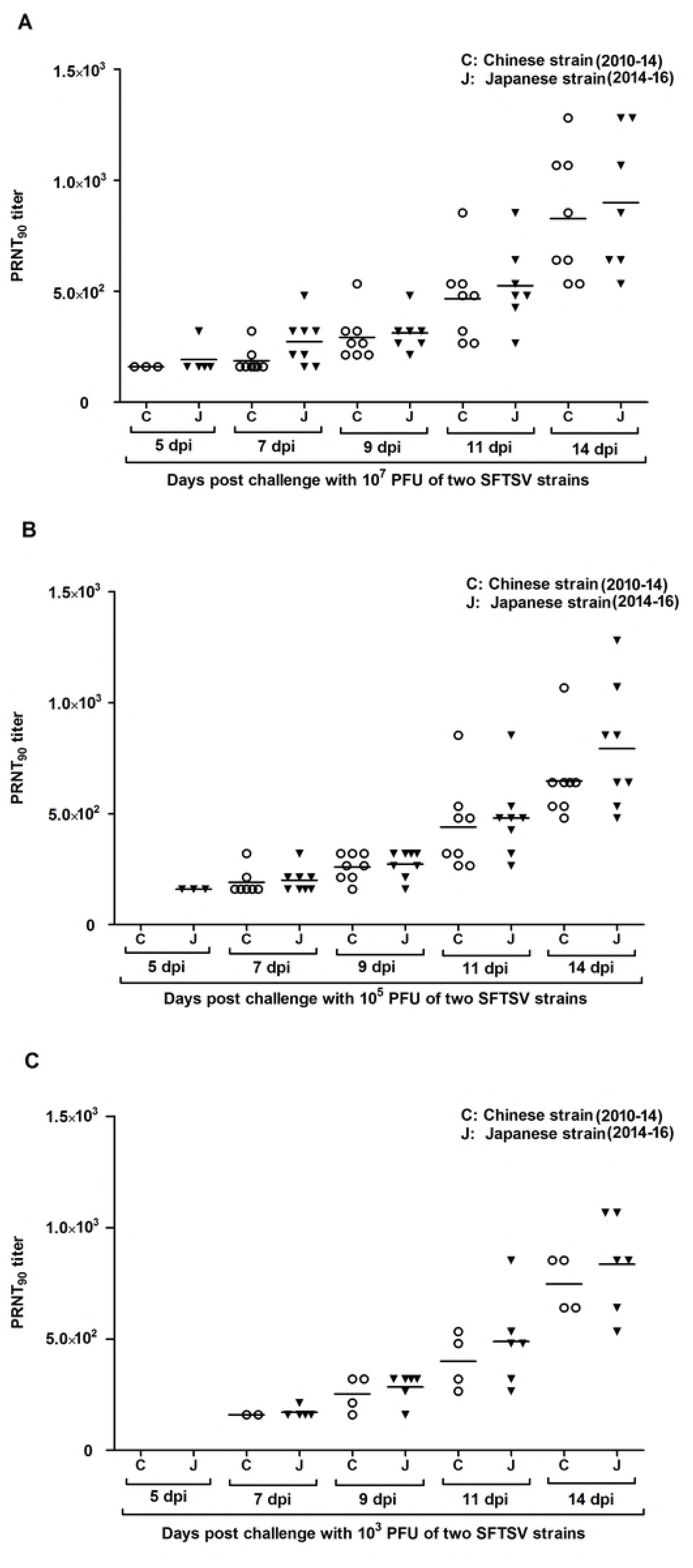
Change in mass of Spotted Doves following SFTSV challenge. Mean change in mass and standard error bars are plotted for daily intervals following challenge for Spotted Doves. Change in mass following SFTSV challenge showed a dose and strain response effect. Birds challenged with 10^7^ plaque-forming units (PFU) of SFTSV strain JS2014 had earlier and greater loss of mass than birds challenged with 10^5^ or 10^3^ PFU of the SFTSV strain. Birds challenged with SFTSV strain JS2014 had greater loss of mass than birds challenged with same dose level of SFTSV strain JS2010.

### Pathogenicity of the SFTSV infection in spotted doves

In the first study, all infected birds underwent a period of anorexia that coincided with detectable viremia in the groups challenged with 10^7^ and 10^5^ PFU of both SFTSV strains. Moreover, a bird challenged with 10^7^ PFU of the strain JS2014 developed symptoms of lethargy, ruffled and eventually died after peak viremia at 7 dpi. In the groups challenged with 10^3^ PFU of both strains, however, only the birds with detectable viremia showed anorexia. No other clinical signs were observed.

A dose-related loss of body weight was detected in birds following challenges with the two SFTSV strains in the first study (Figure 4). The birds challenged with 10^7^ PFU of the strain JS2014 had the largest drop in body mass, an average of 4.7% by 4 dpi. The birds challenged with 10^7^ PFU of SFTSV strain JS2010 showed the greatest body mass loss of 3.6% average at 5 dpi. Challenged with 10^5^ and 10^3^ PFU of the strain JS2014, the birds lost 3.5% and 2.3% of average body mass at 5 and 6 dpi, respectively. Likewise, birds lost 2.4% and 1.3% of average body mass at 6 and 7 dpi, respectively, when challenged with 10^5^ and 10^3^ PFU of the strain JS2010. In comparison, the JS2014 strain appeared to cause the most mass loss earlier and more severe than the JS2010 strain. By 10 dpi the mean mass had either returned to or exceeded its starting level in all challenge groups.

**Figure 4.**
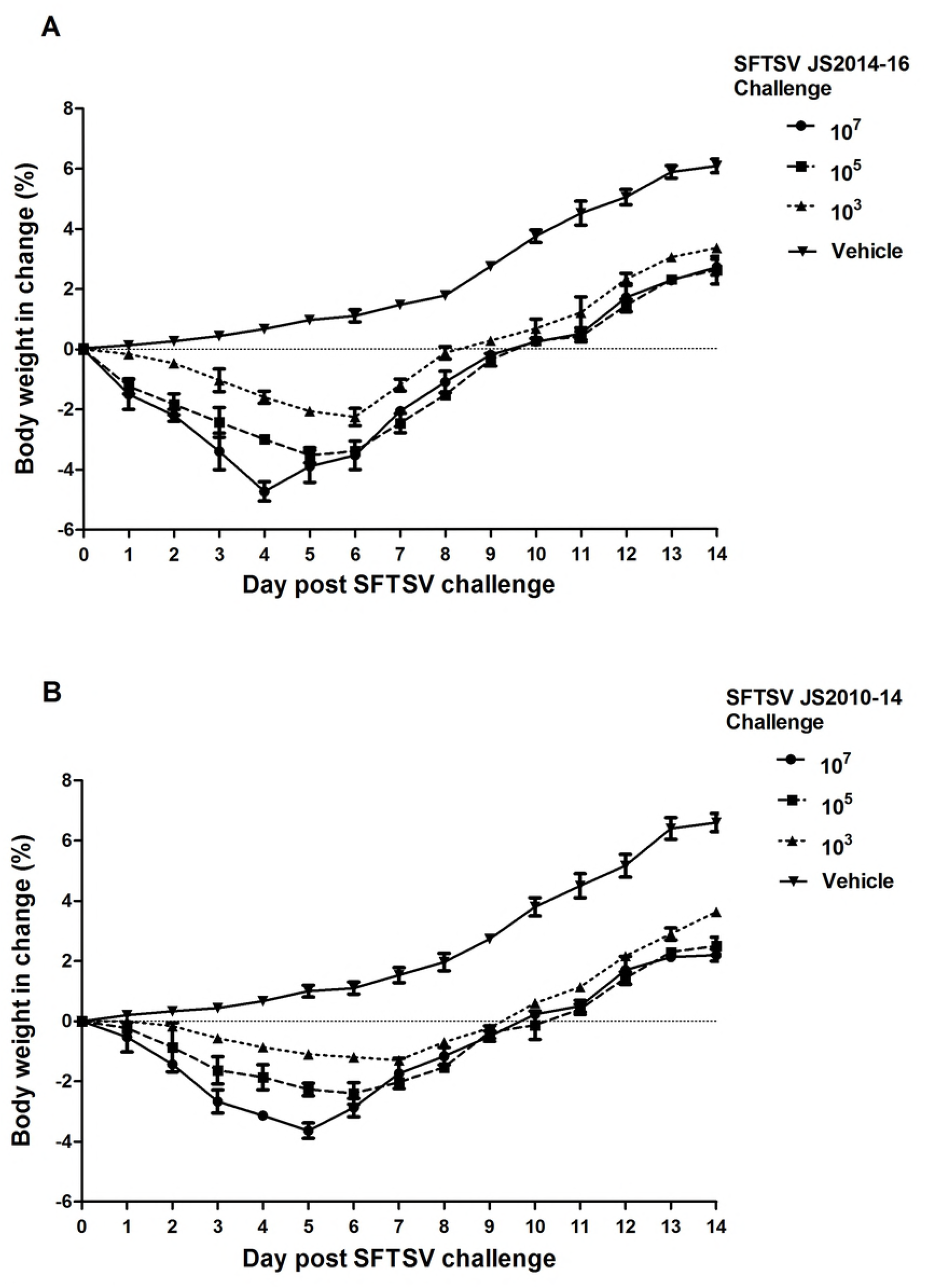
Neutralizing antibody titers of Spotted Doves infected with different challenge levels of two SFTSV strains. Spotted Doves challenged with 10^7^ and 10^5^ PFU of SFTSV (A and B) showed earlier neutralizing antibody than 10^3^ challenge groups (C).

## Discussion

SFTS is an emerging zoonotic disease which can be traced back to initial reports of an unknown infectious disease in rural areas of Hubei and Henan provinces in central China in 2009 [1]. The causative agent was not determined initially due to similar clinical manifestations caused by *Anaplasma phagocytophilum*, Hantaan virus, and *Rickettsia tsutsugamushi* infection [1-2]. An active investigation including viral isolation and molecular characterization was finally implemented resulting in the isolation and confirmation of SFTSV from a farmer in Henan, China [1]. Surveillance data showed that SFTSV has spread to 23 provinces in China from 2010 to 2017 [3-6, 19]. Furthermore, SFTS cases have been reported in other Asian countries including South Korea and Japan [19-21].

Several studies on geographic distribution, genetic diversity, and prevalence of SFTSV genotypes have proposed that there are two major SFTSV genetic lineages, named the Chinese and Japanese lineages [22]. Phylogenetically the SFTSV strains were grouped in 5 clades (A, B, C, D and E) based on the sequences of their genome segments. The clades A, B, C and D were classified ias Chinese lineage, while clade E was classified as the Japanese lineage. SFTSV strains isolated from China fell in all 5 clades, the strains from South Korea were classified into 3 clades (A, D, and E), and all strains from Japan from only clade E [6, 19, 22-23]. At present, the SFTSV strains of clade E were the most widely disseminated in East Asia.

Recent analyses indicate that SFTSV might be originated in the Dabie Mountain area, including Henan, Hubei, Anhui, and Jiangsu provinces, in central China. Several decades ago the virus was transmitted to Shandong Province from Henan Province, and to Liaoning of the Northeastern China and to the Zhoushan Archipelago of China, Jeju Island of South Korea, and Japan from Jiangsu province according to the theory [23]. Transmission in both ways by land or even across the sea could have happened among these areas, mainly of the virus from clade E [23].

Discoveries have been made in the past years about the natural transmission cycle of SFTSV. Previous studies have found that some domestic animals and wildlife can be infected with SFTSV, which might serve as amplifying or reservoir hosts in the natural transmission cycle of SFTSV [10-12, 16]. Our recent study showed that some species of migratory birds, such as swan geese and spotted doves, can both be parasitized by *H. longicornis* and infected by SFTSV naturally [7]. Other studies have reported that migratory bird routes and the distribution of *H. longicornis* in East Asia overlap with the geographic distribution of SFTSV [24-25]. Migratory birds are known to be carriers and transmitters of infectious agents, like the causative agents of influenza, West Nile encephalitis, and Lyme disease [26-27]. Wild birds often travel long distances carrying various parasites, including ticks, which may be infected with viruses and bacteria. It is therefore reasonable to hypothesize that migratory birds may have played an important role in dispersing *H. longicornis-borne* SFTSV in both scenarios, either the birds are infected directly with the virus or the birds are carriers of parasitic ticks that are infected with the virus.

Spotted doves are a common migratory bird in China. In this study, we challenged spotted doves with a Chinese lineage (clade A) and a Japanese lineage (clade E) SFTSV strain to establish a bird laboratory infection model of SFTSV. We were also interested in examining host susceptibility to infection, viral pathogenicity and duration of viremia, if susceptible, in order to assess the potential role of Spotted Doves as a competent host capable of transmitting the virus. The results showed that Spotted Doves were susceptible to both clades of the SFTSV strains. Viremia appeared in all birds challenged with the doses of 10^7^ and 10^5^ PFU of both viral strains. Most of the spotted doves challenged with the dose of 10^3^ PFU were infected and had detectable viremia or SFTSV specific antibodies. Mortality was observed only in the group of the birds challenged at 10^7^ PFU with the clade E virus, or the Japanese strain JS2014 (1/8, 12.5%) at day 7 pi. No birds died in either the 10^5^ or 10^3^ groups of the JS2014 strain and all challenge levels of the Chinese strain JS2010 within the period of the study. This suggests that the infection of SFTSV in spotted doves was primarily self-limiting and infected birds mostly recovered after a period of viremia.

Our data indicate differential severities in the birds by the two clades of the viruses. Of the two viral strains tested, JS2014 led to one out of eight death and higher viremia titers while the birds, inoculated with JS2010, suffered no fatality and had relatively lower virus titers in the blood (Figure 1). We speculate that with higher viremia titers, the birds could transmit the virus to feeding ticks more efficiently. Thus, as a potential amplifying host, spotted doves may be more efficient in transmitting Japanese lineage SFTSV. This is consistent with the studies on the geographic distribution of SFTSV genotypes, i.e., that the SFTSV strains of the Japanese lineage are more widely disseminated igeographically, probably because its higher replication efficacy in migratory birds such as doves.

To date, only a fewer mammals have been used as models for the study of SFTSV infection, including mouse, goat, hamster and macaque [13-16]. To our knowledge our study is the first to use spotted doves as a model for testing the susceptibility of birds to SFTSV infection. The results showed that spotted doves are susceptible to SFTSV, which could be a competent amplifying host for SFTSV, the clade E or the Japanese strains in particular, and may play an important role in long-distance transmission of the virus.

## References

1. Yu XJ, Liang MF, Zhang SY, Liu Y, Li JD, Sun YL, et al. Fever with thrombocytopenia associated with a novel bunyavirus in China. The New England journal of medicine. 2011 Apr 21;364(16):1523-32.

2. Xu B, Liu L, Huang X, Ma H, Zhang Y, Du Y, et al. Metagenomic analysis of fever, thrombocytopenia and leukopenia syndrome (FTLS) in Henan Province, China: discovery of a new bunyavirus. PLoS Pathog. 2011 Nov;7(11):e1002369. doi: 10.1371/journal.ppat.1002369.

3. Robles NJC, Han HJ, Park SJ, Choi YK. Epidemiology of severe fever and thrombocytopenia syndrome virus infection and the need for therapeutics for the prevention. Clin Exp Vaccine Res. 2018 Jan;7(1):43-50. doi: 10.7774/cevr.2018.7.1.43.

4. Ding F, Zhang W, Wang L, Hu W, Soares Magalhaes RJ, Sun H, et al. Epidemiologic features of severe fever with thrombocytopenia syndrome in China, 2011-2012. Clinical infectious diseases : an official publication of the Infectious Diseases Society of America. 2013 Jun;56(11):1682-3.

5. Liu Q, He B, Huang SY, Wei F, Zhu XQ. Severe fever with thrombocytopenia syndrome, an emerging tick-borne zoonosis. The Lancet Infectious diseases. 2014 Aug;14(8):763-72.

6. Hu B, Cai K, Liu M, Li W, Xu J, Qiu F, Zhan J. Laboratory detection and molecular phylogenetic analysis of severe fever with thrombocytopenia syndrome virus in Hubei Province, central China. Arch Virol. 2018 Aug 22. doi: 10.1007/s00705-018-3993-5.

7. Li Z, Bao C, Hu J, et al. Ecology of the tick-borne phlebovirus causing severe fever with thrombocytopenia syndrome in an endemic area of China. PLoS Negl Trop Dis 2016;10: e0004574.

8. Wang S, Li J, Niu G, Wang X, Ding S, Jiang X, et al. SFTS virus in ticks in an endemic area of China. The American journal of tropical medicine and hygiene. 2015 Apr;92(4):684-9.

9. Yun SM, Lee WG, Ryou J, Yang SC, Park SW, Roh JY, et al. Severe fever with thrombocytopenia syndrome virus in ticks collected from humans, South Korea, 2013. Emerging infectious diseases. 2014 Aug;20(8):1358-61.

10. Li Z, Hu J, Bao C, et al. Seroprevalence of antibodies against SFTS virus infection in farmers and animals, Jiangsu, China. J Clin Virol 2014;60:185-9.

11. Ding S, Yin H, Xu X, Liu G, Jiang S, Wang W, et al. A cross-sectional survey of severe fever with thrombocytopenia syndrome virus infection of domestic animals in Laizhou City, Shandong Province, China. Japanese journal of infectious diseases. 2014;67(1):1-4.

12. Niu G, Li J, Liang M, Jiang X, Jiang M, Yin H, et al. Severe fever with thrombocytopenia syndrome virus among domesticated animals, China. Emerging infectious diseases. 2013 May;19(5):756-63.

13. Chen XP, Cong ML, Li MH, et al. Infection and pathogenesis of Huaiyangshan virus (a novel tick-borne bunyavirus) in laboratory rodents. J Gen Virol 2012;93:1288-93.

14. Jin C, Liang M, Ning J, et al. Pathogenesis of emerging severe fever with thrombocytopenia syndrome virus in C57/BL6 mouse model. Proc Natl Acad Sci U S A 2012;109:10053-8.

15. Jin C, Jiang H, Liang M, et al. SFTS virus infection in nonhuman primates. J Infect Dis 2015;211:915-25.

16. Jiao Y, Qi X, Liu D, et al. Experimental and natural infections of goats with severe fever with thrombocytopenia syndrome virus: evidence for ticks as viral vector. PLoS Negl Trop Dis 2015;9:e0004092.

17. Lapointe DA, Hofmeister EK, Atkinson CT, Porter RE, Dusek RJ. Experimental infection of Hawai‘i’Amakihi (hemignathus virens) with West Nile virus and competence of a co-occurring vector, culex quinquefasciatus: potential impacts on endemic Hawaiian avifauna. J Wildl Dis. 2009 Apr;45(2):257-71.

18. Li Z, Cui L, Zhou M, Qi X, Bao C, Hu J, et al. Development and application of a one-step real-time RT-PCR using a minor-groove-binding probe for the detection of a novel bunyavirus in clinical specimens. Journal of medical virology. 2013 Feb;85(2):370-7.

19. Li Z, Hu J, Cui L, Hong Y, Liu J, Li P, et al. Increased Prevalence of Severe Fever with Thrombocytopenia Syndrome in Eastern China Clustered with Multiple Genotypes and Reasserted Virus during 2010-2015. Sci Rep. 2017 Jul 26;7(1):6503. doi: 10.1038/s41598-017-06853-1.

20. Kim KH, Yi J, Kim G, Choi SJ, Jun KI, Kim NH, et al. Severe fever with thrombocytopenia syndrome, South Korea, 2012. Emerging infectious diseases. 2013 Nov;19(11):1892-4.

21. Takahashi T, Maeda K, Suzuki T, Ishido A, Shigeoka T, Tominaga T, et al. The first identification and retrospective study of Severe Fever with Thrombocytopenia Syndrome in Japan. The Journal of infectious diseases. 2014 Mar;209(6):816-27.

22. Yoshikawa T, Shimojima M, Fukushi S, Tani H, Fukuma A, Taniguchi S, et al. Phylogenetic and Geographic Relationships of Severe Fever With Thrombocytopenia Syndrome Virus in China, South Korea, and Japan. The Journal of infectious diseases. 2015 Sep 15;212(6):889-98.

23. Liu JW, Zhao L, Luo LM, Liu MM, Sun Y, Su X, Yu XJ. Molecular Evolution and Spatial Transmission of Severe Fever with Thrombocytopenia Syndrome Virus Based on Complete Genome Sequences. PLoS One. 2016 Mar 21;11(3):e0151677. doi: 10.1371/journal.pone.0151677. eCollection 2016.

24. Yun Y, Heo ST, Kim G, Hewson R, Kim H, Park D, et al. Phylogenetic Analysis of Severe Fever with Thrombocytopenia Syndrome Virus in South Korea and Migratory Bird Routes Between China, South Korea, and Japan. Am J Trop Med Hyg. 2015 Sep;93(3):468-74. doi: 10.4269/ajtmh.15-0047.

25. Shi J, Hu S, Liu X, et al. Migration, recombination, and reassortment are involved in the evolution of severe fever with thrombocytopenia syndrome bunyavirus. Infect Genet Evol 2017;47:109-17.

26. Reed KD, Meece JK, Henkel JS, Shukla SK. Birds, migration and emerging zoonoses: west nile virus, lyme disease, influenza A and enteropathogens. Clin Med Res 2003;1:5-12.

27. Lee KH, Medlock JM, Heo ST. Severe fever with thrombocytopenia syndrome virus, Crimean-Congo haemorrhagic fever virus, and migratory birds. J Bacteriol Virol 2013;43: 235-43.

28. Fan Z, Liu D, Tang X, Huang B, Qiu J, Qiu B, et al. A surveillance of avian influenza virus infectionin wild birds. Chinese Journal of Preventive Veterinary Medicine. 2012, 34(2):96-99.

